# The Import of Skin Tissue Dynamics in Tactile Sensing

**DOI:** 10.1101/2021.09.06.459180

**Authors:** Udaya B. Rongala, Andre Seyfarth, Vincent Hayward, Henrik Jörntell

## Abstract

The mammalian skin is a densely innervated soft tissue where embedded neural mechanoreceptors report to the brain the mechanical events arising at its surface. Up to now, models of the transformations of these events into neural signals relied on quasistatic or viscoelastic mechanical models of the skin. Here, we employed a model which, in addition to elasticity and viscosity, accounted for mass to accurately reproduce the propagation of mechanical waves observed in vivo. Skin dynamics converted sensory inputs into rapidly evolving spatiotemporal patterns that magnified the information made available to a population of mechanoreceptors. Accounting for dynamics greatly enhanced the separability of tactile inputs and was efficient for a large range of mechanical parameter values. This advantage vanished when these parameters were set to approximate the quasistatic or viscoelastic cases.

---

Barring inputs from claws, nails, hoofs, scales, horns, or fins, in most animals touch begins in the skin. The glabrous mammalian skin — a type of soft tissue enabling manipulation or locomotion — is by far the most studied touch sensitive tissue. Mechanical models of glabrous skin have been proposed to clarify the manner in which deformations are transformed into signals available to mechanoreceptors located beneath the surface. Skin tissue is also often indirectly stimulated through intermediaries such as hairs, whiskers, feathers, tools or clothing interposed between touched objects and sensitive tissues. The importance of the dynamic properties of the skin investigated in present study would, with adjustments, nevertheless also extend to these cases.

Extant models of the transformation between skin deformation and mechanoreceptor response typically appeal to the mechanics of continuum media to describe how variations in surface boundary conditions translate into a strain field below the surface (1–9). These models have been effective for producing predictions in the quasi-static and viscoelastic cases where mass properties do not enter in the model formulation. In other words, these models assume that elasticity and viscosity dominate over inertial contributions to tissue behaviour. Several recent studies, however, have shown that mechanical waves propagating in the soft tissues of the extremities possess behavioural importance during tactile interaction (10–15). It follows from these studies that quasi-static and viscoelastic models of the skin furnish an incomplete picture of the process of transformation from surface boundary conditions to signals available to the skin mechanoreceptors. The most advanced model of transduction of mechanical inputs into neural messages, to date, considers a simplified delay model of mechanical wave propagation in the skin (16).

Here, we advance the hypothesis that the mass-induced dynamic properties of the skin, beyond viscoelasticity, magnify the amount of information potentially available to the somatosensory system through a diversification of spatiotemporal activation patterns among a population of mechanoreceptors. To put this hypothesis to the test, we employed a model that accurately captures how brief inputs determine the propagation of mechanical waves in the skin (Materials and Methods). This model was used to evaluate tactile information available to the mechanoreceptors when the group velocity of wave propagation and its decay characteristic time were set to approximate that of biological skin. To this end, we quantified the diversity of the signals made available to a population of mechanoreceptor through a principal component analysis and computed the total entropy of these signals for standardised brief inputs. To augment the biological plausibility of our model, we also evaluated these measures when the mechanoreceptors were modelled as spiking units.

## Model captured biological skin dynamics

To rest our model on empirical data, we performed *in vivo* measurements of the mechanical evoked response to inputs localised in time and in space to reveal the dominant dynamic properties of the human fingertip considered as a propagation medium. The evoked response was relevant to the neural message collected in the extremities through two main quantities: wave group velocity and amplitude decay time. Group velocity, which characterises the velocity at which information is transported in media, affects the relative timing within the neural messages made available to the brain. Decay, which depends on the size and shape of the domain in which waves develop and on the properties of the propagation medium, affects how much information about dynamic events is made available to mechanotransduction processes over behaviourally relevant time scales.

Actual skin is a dispersive poroelastic medium that exhibits complex behaviours that depend on biological factors which are difficult to disentangle, *e.g*. (17, 18). Because steady sine waves or superposition thereof do not carry information, we preferred using brief test inputs in the form of displacement Gaussian wavelets, 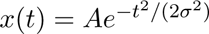, with fixed amplitude, *A*, and characteristic time, *σ*, ranging from 1.0 ms to 0.2 ms (Materials and Methods), Fig. 1A. These time scales correspond to behaviourally relevant frequencies in the range from 160 to 800 Hz (Materials and Methods) during active touch (19–21). From the superposition principle (22), complex inputs can be approximated by weighted combinations of such Gaussian wavelets, despite their lacking orthogonality (23).

**Figure 1:**
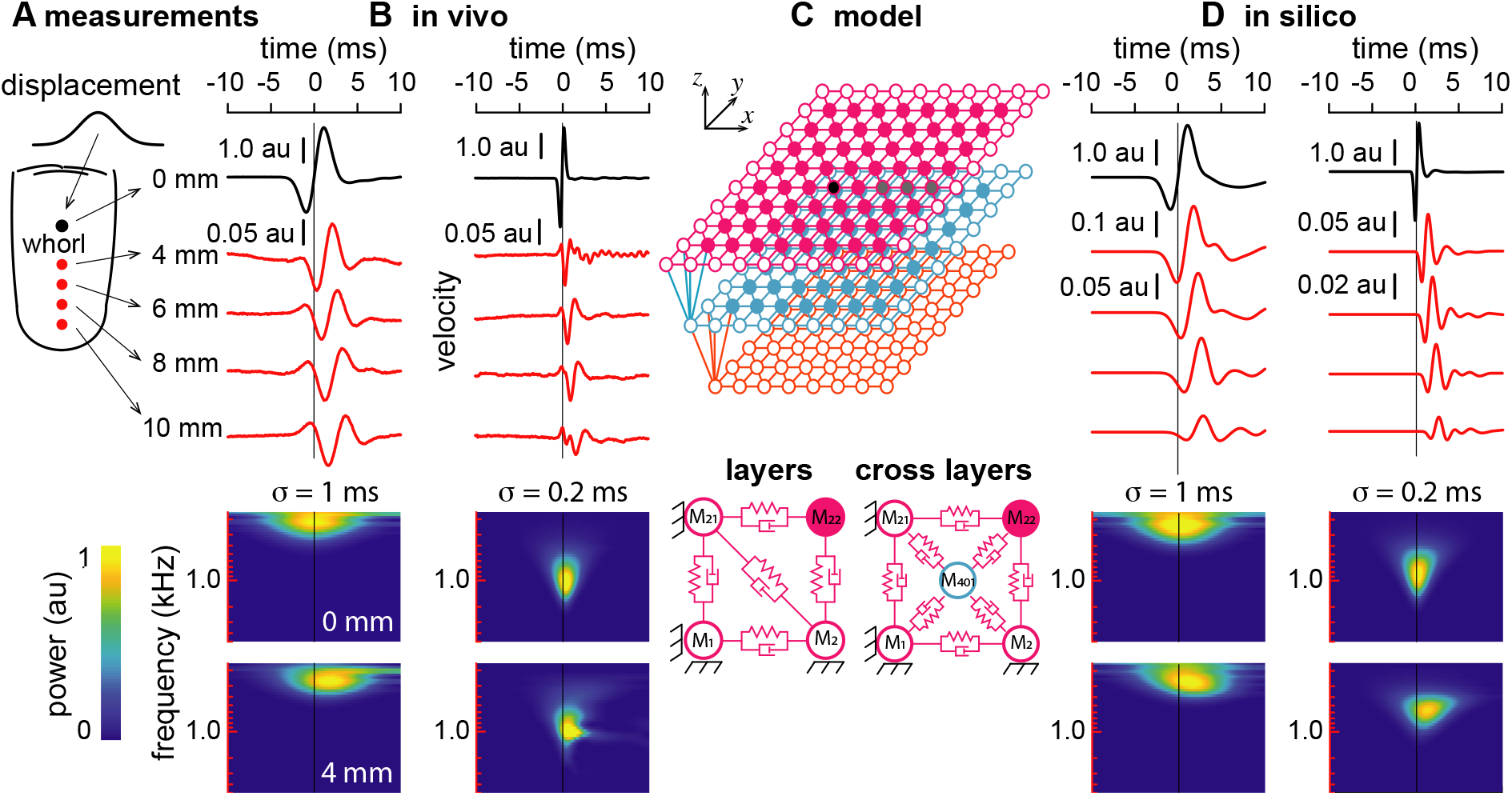
Damped mass-spring model captures skin dynamics. (**A**) *In vivo* measurements of skin dynamics (Materials and Methods). Gaussian wavelet of surface normal displacement centred at time *t* = 0 applied to the index finger whorl by a 1.0 mm circular punch. Velocity measurements at five locations distant from 0.0, 4.0, 6.0, 8.0, and 10.0 mm. (**B**) Evoked responses (red lines) in the normal direction for characteristic times *σ* = 1.0 ms and *σ* = 0.2 ms. Spectrograms of the measured velocity at 0 mm and 4 mm away from the locus of stimulation (Materials and Methods). (**C**) Three-layer damped mass-spring skin model. Masses, shown as nodes, were connected by springs and dampers represented as links of 1.0 mm. Actual model comprised 509 nodes, the top layer comprising 14×14 nodes. Nodes shown as circles were fixed nodes to establish a boundary condition. The tuned parameters were the stiffness and the damping of the links, *K* = 2.5 10 N/m and *C* = 5 × 10^−4^ Ns/m respectively, the individual masses were *M* = 1.0 mg for a total of 0.5 g of simulated tissue. Spring-dampers between masses within a same layer and across different layers. (**D**) Model responses at different distances from the locus of stimulation for the same input wavelets. Continuous wavelet transform extracted the signal power over mechanoreceptors activity (Materials and Methods).

The test inputs were applied at the centre on the fingerprint whorl of one male participant’s left index finger, see Fig. 1A. Responses were collected at distances of 4, 6, 8, and 10 mm. Attenuation was rapid near the locus of stimulation and slower after a few millimetres indicating that, in humans, hundreds, if not thousands of mechanoreceptors, would be stimulated by such inputs. This pattern was consistent with the propagation of mechanical waves in inertio-visco-elastic half spaces and may have corresponded to near- and far-field solutions (24) for both slower and faster wavelets shown in Fig. 1B.

The model (Materials and Methods) comprised three layers of 14×14 nodes, 13×13 nodes, and 12×12 nodes respectively, see Fig. 1C, for a total of 509 nodes connected in a triangulated network within layers. The distance between nodes was 1.0 mm. Generic nodes in a layer were connected to six neighbours, boundary nodes to three neighbours. Nodes within a layer were connected to four neighbours of the layer below, resulting in a dense pyramidal network. The deepest layer was connected to fixed boundary nodes via dampers. The model responses could be made to match the in vivo responses, as shown in Fig. 1D. After tuning, the characteristic time was about 3.0 ms and the group velocity was about 1.7 m/s. The model reproduced the overall dispersive character of skin-borne mechanical waves, albeit not completely accurately.

## Dynamic skin model generated rich information from brief stimuli

The relationship between skin deformation and mechanoreceptor sensory signalling is a debated issue. Earlier models struggle to explain empirical neural responses to components of the strain tensor field (there are six components for small deformations) or resort to correlations (1, 2, 25, 6, 8). These difficulties probably originate from an unsupported assumption that the skin can be modelled as a homogeneous and continuous material. The skin is in fact made made of fibrous, cellular, and bi-phasic tissues. At the length scale of mechanoreceptors, the modes of deformation would fail to follow the rules of continuum mechanics (26). Even if correlations with components of a modelled tensor field can be found, it remains that mechanoreceptors respond to tissue deformation in ways that are largely unknown, let alone in the dynamic case. For example, Meissner corpuscles of the mammalian digital skin, which are encapsulated Schwann cell helices about 10 μm in diameter connected to surrounding cells by randomly oriented collagen fibres networks (27), fail to exhibit clear selectively to any particular mode of deformation associated to invariants of the Cauchy-Green deformation tensor. At the scale of membranes, mechanotransduction was found to appel to structural mechanics that cannot be reduced to the averaging components of a deformation tensor (28). This point is further discussed in Section “Effect of spike signalling”. Nevertheless, *in vivo* studies consistently demonstrate high temporal resolution across the whole diversity of skin mechanoreceptors (29–34).

The model adopted inter-node distance, *i.e*. strain, as the best available proxy for representing the activation of mechanoreceptors since it represented the temporal evolution of the information made available by skin dynamics. A principal component analysis (PCA) across the population of simulated mechanoreceptors assessed this information. Figure 2 shows mechanoreceptor activity for four different inputs, indent, push, quad, and slide along with the evolution of the corresponding principal components. Figure 2A shows mechanoreceptor activity of proximal and distal nodes for input indent where a single node is displaced in the normal direction. The signal was modified as a result of transformations arising from wave propagation. With input push, Fig. 2B, four neighbouring nodes were simultaneously stimulated. The first two components initially evolved in manner similar to that of input indent. Separability of the input signals, however, was ensured by the next components which evolved differently. Input quad, Fig. 2C, where four nodes were stimulated in different directions, led to distinctively different evolutions of all the principal components. With input slide where displacements are in a lateral direction, Fig. 2D, the first component reflected the ensemble response of a population of mechanoreceptors. The first component was distinctive from that of the other cases, with separability further enhanced by the higher order principal components. The ability to separate different inputs can be further exemplified when comparing the evolution of some of the first principal components, for instance, PC 1, PC 5, and PC 10 as shown in Fig. 2E.

**Figure 2:**
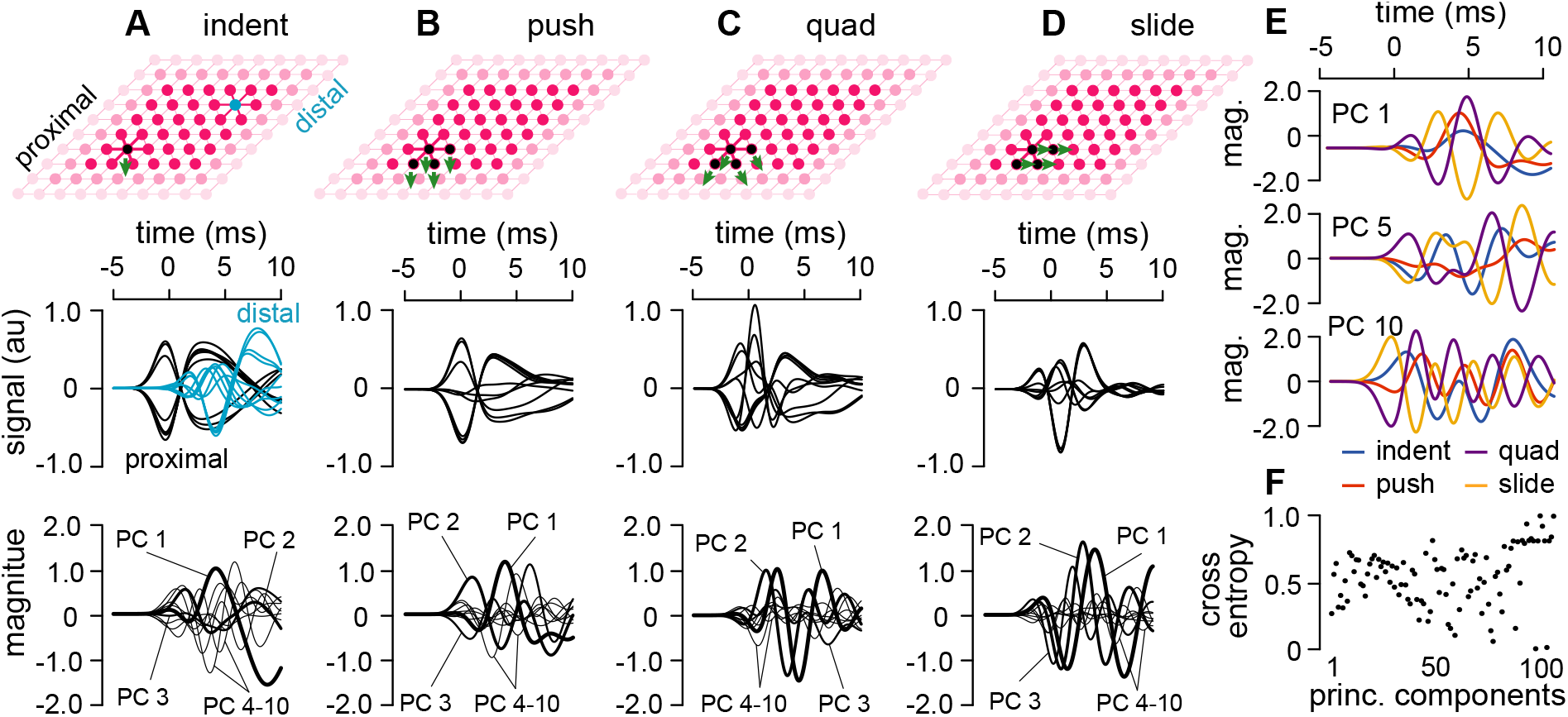
Effect of skin dynamics. (**A**) Input indent was a Gaussian wavelet of normal indentation with *σ* = 1.0 ms characteristic time applied to the node in black. Mechanoreceptor outputs (black lines) from links connected to the activated node and mechanoreceptor outputs from ten distal links (blue lines). First ten principal components (first: thick line, second: thinner line, and so on). (**B**) Input push involved four neighbouring nodes stimulated in the normal direction (green arrows). Mechanoreceptor outputs from ten links connected to the activated nodes and corresponding first ten principal components. (**C**) Input quad stimulated four nodes in four different directions. (**D**) Input slide stimulated four neighbouring nodes in a lateral direction. Mechanoreceptor outputs from ten links directly connected to the activated nodes and ten first principal components. (**E**) Principal components 1, 5, and 10 for four different inputs. (**F**) Entropy across four different inputs (cross entropy) for principal components 1 to 100.

## Available information

Individual principal components best represented activity distributions across a population of mechanoreceptors given particular inputs (Figs. 2A–D). The variance explained by any particular principal component here reflected the significance of any specific activity distribution across the mechanoreceptor population. The magnitude of the first principal component, Fig. 2E, accounted for the greatest part of the variance. In a system with many degrees of freedom, signals can be decomposed into many possible activity distributions, resulting in a high number of principal components and enabling separability of many input signals. For example, the simple input of Fig. 2A required fifteen components to explain 95% of the variance and thirty components to explain 99.9% of it. To assess differences across conditions the evolution of a few of the first principal components for four different inputs is shown in Fig. 2E. While the first few components might suffice for an efficient separability of the different inputs, the higher order components enhanced separability. To further quantify the separability of the different inputs, we calculated the cross entropy for each individual principal component Fig. 2E. While the principal components of higher order may have accounted individually for a small part of the total response, they represented information that could further increase input separability.

## Available information is reduced in the absence of mass properties

Different species, different individuals, different skin areas within an individual have different dynamics. Would certain skin properties be beneficial to promote the advantage owed to dynamical effects? In order to assess the impact of skin dynamics, we generated an instantiation of our model by eliminating the inertial term thus reducing it to a viscoelastic model (Materials and Methods). The dynamics was largely lost as shown in Fig, 3A. The responses of proximal mechanoreceptors sensors reflected the temporal profile of the indentation, in contrast with the responses evoked in the dynamic skin model. Compare Fig. 2A with Fig. 3A for the same input where in the latter case, as it should, the activity the distal nodes vanished. To quantify differences in the amount of information available by each skin model setting, we calculated to entropy across the principal components for each given input.

**Figure 3:**
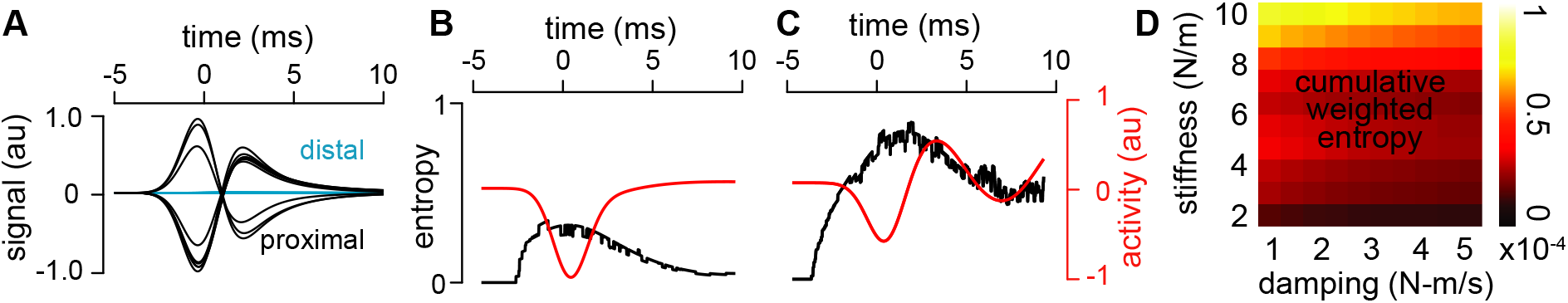
Information loss in the viscoelastic case quantified by entropy. (**A**) Mechanoreceptor outputs when the skin model was viscoelastic. (**B**) Weighted entropy associated with the principal components of the outputs of the mechanoreceptor population for the indentation stimulus for a given input in the viscoelastic case (Fig. 2A). (**C**) Same quantities with skin parameters tuned to resemble actual skin’s dynamics (Figs. 1B and 1E). (**D**) Cumulative entropy as a function of stiffness and damping model parameters (Materials and Methods).

The loss of available information was evident when comparing Fig. 3B corresponding to the viscoelastic case with Fig. 3C which shows the case when the parameters were set to approximate actual skin. In the latter case, the entropy remained high beyond the stimulation activation peak. The distribution of activity across the mechanoreceptors varied during and after the stimulus, which would enable a decoder not only to report when an input occurred but also to discriminate between its different phases: onset, peak, and decay. The entropy, Fig. 3C (black line), indicating the possibility to separate inputs, was indicative that a subset of the principal components were already active during the early period of the time course of the stimulation. The same was true for later parts of the response.

Figure 3D shows that for most parameter settings, the cumulative entropy (area under the curve) was not greatly altered by a modification of the mechanical parameters, except when stiffness was very low or very high. These data shows that high stiffness was associated high entropy. The low dependency on damping suggests that this parameter mostly impacts sensing temporal resolution. A noise-free model, however, would leave entropy invariant under changes in signal magnitude, which is unrealistic (35).

## Effect of spike signalling and noise

Actual mechanoreceptors are noisy, leading to uncertainty. Some of the uncertainty is owed to the discrete nature of spiking outputs, since discretisation is always associated with loss of information, and some comes from the biophysics of transduction, even if noisy properties may come with advantages from a biological perspective (36, 37).

To illustrate the impact of noise, first consider the hypothetical noise-free responses of ten simulated mechanoreceptors to a brief indentation stimulus, Fig. 4A. Because a neural spiking can only signal positive values, the model ignored the negative part of the sensor signal. This model is in line with directional selectivity observed among fingertip sensors in humans (38) and is justified by the fact that the molecular-scale mechanics that underlies the opening of the mechanotransduction channels (39) is not well understood as it involves a molecular-scale kinematics driven by the deformation of the embedding skin tissue (40). Studies from other organs where mechanotransduction takes place show that normal and shear strain sensing depend on the configuration of the coupling (41–43).

**Figure 4:**
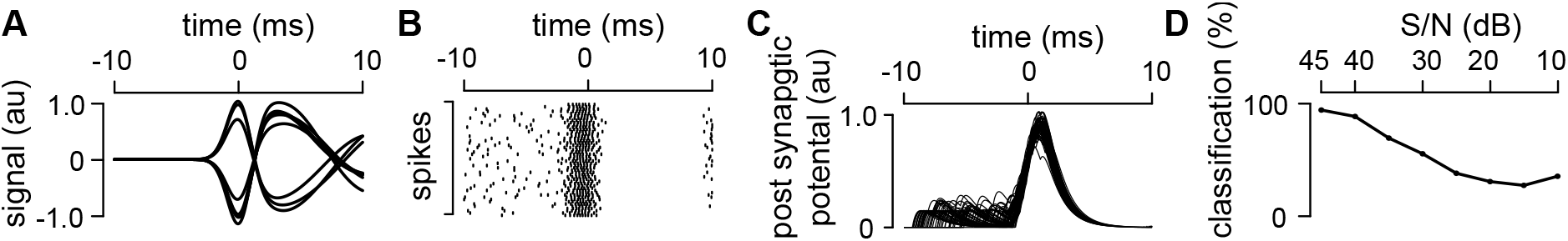
Sensitivity to spike coding. (**A**) Noise-free, mechanoreceptor responses for the indent stimulus (Fig. 2A). (**B**) Raster plot of the spike outputs simulated by the stochastic Izhikevich neuron model (25 dB SNR, Materials and Methods) across fifty repetitions. (**C**) Fifty overlaid spike train responses convoluted using a kernel function (Material and Methods). Fifty overlaid spike train responses convoluted using a kernel function (Material and Methods). (**D**) Classification performance across signal-to-noise values.

Then, compare the noise-free signals to the spike responses signalled by simulated primary sensory neurones for the same input during fifty repetitions as in Fig. 4B (Material and Methods). Owing to discretisation, the spiking output represents a lossy transformation of the underlying mechanoreceptor signal where the average inter-spikes interval reflects the amount of excitation in the membrane potential.

Our model, by simulating postsynaptic responses to fifty repetitions of the same input, see Fig. 4C, represented the information made available by the sensory neurones to the second-order neurones, i.e. the cuneate nucleus output neurones (Material and Methods). To evaluate information loss associated with the stochasticity of the transduction, the performance of a standard Support Vector Machine classifier (Material and Methods) was evaluated as a function of the noise injected in the model. Figure 4D shows how input separability was impaired as the signal-to-noise ratio of the neuronal spike generating mechanism decreased.

## Summary and Discussion

Using a model of wave propagation in the skin (Materials and Methods), we showed that for brief, localised stimuli used as model interactions, this dynamics profoundly impacted on the information available to mechanotransduction. These elementary stimuli support general representations of complex, distributed stimuli by virtue of the principle of superposition (Material and Methods).

The dramatic increase of information available to neuro-transduction arose from a combination of effects. The low group velocity of tactile waves combined with a slow rate of decay, implies that any mechanical event localised in time and in space could stimulate a considerable population of mechanoreceptors. In the human fingertip, for example, this velocity varies around 3.0 m/s (44–46) and the average distance between mechanoreceptors is about 100 μm (47). In a 3.0 mm neighbourhood, a sub-population of mechanoreceptors would experience the consequences of brief stimuli with differential delays up to about 1.0 ms, which is within their temporal resolution (29, 33), reducing the correlation among the responses of a population of mechanoreceptors. This effect was reflected in the results reported in Fig. 2E where the consequences of the dynamics was seen in the many principal components required to represent a particular stimulus. These many components translate into high separability of similar stimuli such as those exemplified in Fig. 2.

For complex stimuli, as in Fig. 2B,C,D, an effect of wave propagation is the creation of interferences that increase the complexity of signals made available to a population of receptors sampling the medium. This effect in visible in Fig. 2F where the entropy of the signals made available by a population of mechanoreceptors is high even for lower order principal components. In biology, there must also be a trade-off between the density of the mechanoreceptor distribution in the skin and the brain’s available computing capacity, which may have been optimised through evolution. Mechanoreceptor distributions are known to be highly variable across both body regions and across species (48).

Adaptation, thresholding, and averaging are noise-reducing strategies documented in neural sensory systems. A novel possibility afforded by pre-neural, dynamic connections within populations of mechanoreceptors for complex stimuli, as in Figs. 2, is combating noise. In contrast, a population of mechanoreceptors activated independently are akin to a collection of independent sources of signal polluted by independent noises. Such configuration procures no advantage, except in the rare occasions when all the sources transmit the same signal. Conversely, if the sources share some regularities, induced here by skin dynamics, then efficient noise reduction strategies become possible. Broad similarities could be drawn in the auditory domain (49). In vision, this effect could be related to minute eye movements that are thought combating noise by accentuating the high spatial frequency harmonics in stimuli (50).

The diversity of mechanical properties of actual skins begs the question of its impact on behaviour. It can be observed that different animals have different skins but little data on their tactile acumen is available besides behavioural observations (51, 52, “Moles can touch a small prey item … and take the prey into the mouth, all in about 400 ms”; “Clean, dirty, wet, and dry foods are all dunked with similar frequency. Raccoons probably handle their food in water to provide greater tactile sensation”.). To our knowledge, there is no study with humans that links directly tactile acumen to the mechanical properties of the skin. If we accept that skin hydration may be used as a proxy for skin elasticity, then such links exist. It was observed that high levels of hydration did not affect detection thresholds to vibrotactile stimuli (17), which is consistent with the fact that steady vibrations do not contain information, however, skin hydration altered the perception of textured surfaces which our findings may contribute to explain (17, 53). It was shown that the pleasantness of surfaces as well as material discrimination performance were also strongly correlated with skin hydration (21, 54).

Our model has applications in robotic artificial sensing which to date has been addressed mostly as a quasi-static or viscoelastic problem. It has also applications in the design of sensory compensation and supplementation devices for neuroprostheses and in the elucidation of early processes in biological touch.

## Materials and Methods

### Apparatus

*In vivo* skin dynamics were investigated using an in-house constructed apparatus arranged as indicated in Fig. 5A. The index finger was resting in a supinated posture upon a stabilising cradle. A one millimetre circular punch impinged on the skin through an electrodynamic motor (55) reconfigured to operate as moving magnet machine. The over-moulded housing holding the voice coils were rigidly connected to an identical motor which was driven in phase opposition to annihilate the total mechanical momentum. This tandem configuration eliminated the perturbations that could be transmitted to the resting finger by the structure. To further attenuate unwanted signals, the motor tandem was supported by a frame through a viscoelastic suspension. The motors were powered through a load balancing circuit by a class-D audio amplifier (Model AA-AB31184, Wondom, Nanjing, PRC).

**Figure 5:**
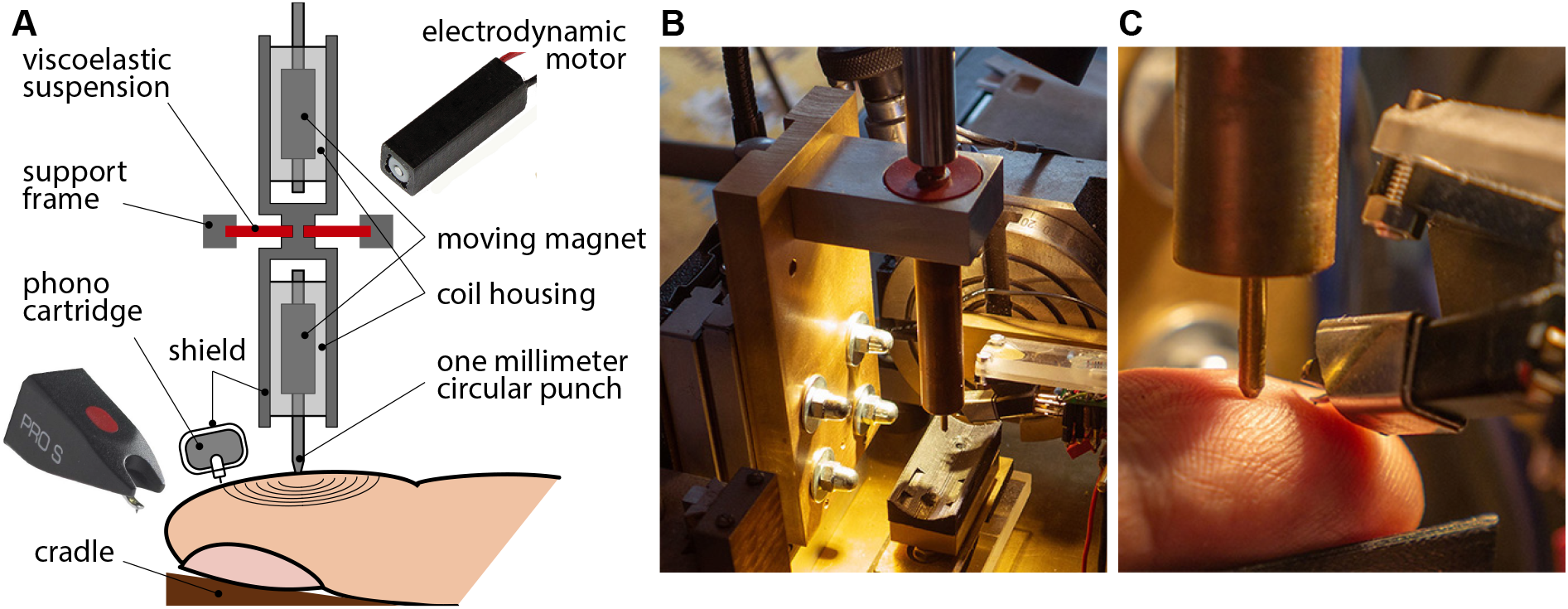
Apparatus. (**A**) General arrangement, see text. (**B**) General view. (**C**) Close up.

A phono cartridge (OM Pro S, spherical stylus, Ortofon, Nakskov, Denmark, sensitivity 10 Vs/m) measured the velocity of the skin displacements. The cartridge was shielded by ferromagnetic foil from magnetic disturbances. The signal was amplified by 20 dB by a custom-made, flat response, low noise preamplifier before being logged at 44 kHz with 16-bit resolution by an audio signal processor (Bela mini, Augmented Instruments Ltd, London, UK). The probe was micro-positioned in x-y-z space and the sensor micro-oriented around two axes about a point fixed coinciding with its tip, Fig. 5B, such that skin displacements could be measured in finger-surface coordinates, Fig. 5C.

### Stimuli

The stimuli were displacement Gaussian wavelets, *Ag*(*t*) where 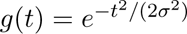, which are frequently used to represent brief interactions between solids. For convenience the wavelets were centred at zero. The corresponding velocity Gaussian wavelet is −*t*/*σ*^2^*g*(*t*). Stimuli had a displacement amplitude of about 0.2 mm which was the same for all characteristic times. The transfer function of the motor tandem was that of a second order mechanical oscillator with a natural frequency of 70 Hz and a damping ratio of 0.2. After inverse filtering giving the system a critically-damped acceleration response, the bandwidth was extended by one octave below the natural frequency. The Fourier transform of a Gaussian wavelet displacement is 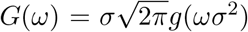 and that of the velocity is *iωG*(*ω*) which for *σ* = 1.0 ms peaks at 160 Hz. Signal energy drops rapidly on either sides of this frequency (*σ* = 0.2 ms and 800 Hz, respectively). Stimuli were accurately produced (Fig. 1B,D).

Broadband tribological measurements (56) agree with the view that the sliding of the human fingertip, which is a multi-scale surface spanning length scales from 10 nm (corneocytes) to 100 μm (ridges) (57), on another rough surface, i.e. a texture, can be modelled by the superposition of random micro-collisions at many different intensities and time scales. This view was empirically vindicated in the case of the largest length scales in dry and wet contacts (58, 59). Each of these micro-collisions can be represented by Gaussian wavelets of different amplitudes and characteristic times.

### Mechanical model

The soft tissues of sensitive extremities in mammals possess complex mechanical properties, however, over short distances, computationally efficient mass-spring models can provide a good representation of the effects of mechanical wave propagation. Spring-mass models of solids have met with success in a variety of domains such as structured materials (60, 61), biomechanics (62, 63), seismology (64), acoustics (65), computer graphics (66–68), virtual reality (69), *inter alia* because of key advantages, chief among them are computational efficiency and ease of tuning to match empirical data. Disadvantages of spring-mass models include lack of observance of fundamental invariants such as those of the Cauchy-Green deformation tensor which precludes using spring-mass models to correctly represent shear or variations of volume. Another shortcoming is lack of continuity, which finite element, boundary elements, or meshless methods (70) can provide. However, the latter methods face challenges when representing dynamic phenomena such as waves (71, 72) whereas spring-mass models accommodate the simulation of wave propagation in solid media quite naturally.

The mass-spring-damper skin model was organised in three layers of concentrated masses connected by springs and dampers as described in Fig. 1C, caption, and surrounding text. Such systems are governed by a system of linear, coupled, second-order ordinary differential equations which is often written,

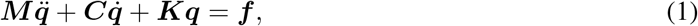

where ***M***, ***C*** and ***K*** are symmetric matrices representing mass, stiffness, damping (Supplementary Material) of the system, ***f*** is the forcing input vector, and ***q*** the vector representing the displacements of the nodes in three dimensions. If there are *N* nodes, then dim ***q*** = dim ***f*** = 3*N*. The system possesses 6*N* states and can be expressed is state-space form, 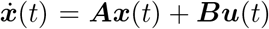, where ***x*** is the state vector, ***A*** the state transition matrix, ***B*** the input matrix, and ***u*** the input. Posing 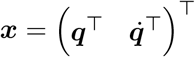 and *u* = (0^⊤^ *f*^⊤^)^⊤^, Eq. (1) can be rewritten (73),

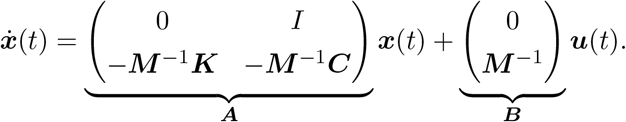

For the viscoelastic case, the model was,

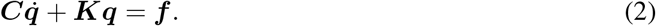

The above systems were solved numerically using the fifth order Runge-Kutta method (`ode45` solver, MatLab^™^). The nominal parameters were set as indicated in the main text.

### Frequency Analysis

Frequency decomposition of the measured and simulated signals (Fig. 1) were performed by continuous wavelet transform (function `cwt`, MatLab^™^).

### Data analysis

Principal component analysis was performed on the sensory responses (Fig. 2, Fig. 3A), (function pca, MatLab^™^). In addition to the magnitude of the principal components, the variance of the responses for each principal component explained was computed. The higher the number of principal components to explain a given level of variance, the higher the diversity of the sensory responses.

Cross entropy (Fig. 2F) was calculated per principal component. Entropy, *H*_n_, was calculated across all the principal components for each time-step *n*. Weighted entropy (Fig. 3B,C) was the entropy measure multiplied by the total sensory response within the time window (function `trapz`, MatLab™). Cumulative entropy (Fig. 3D) was calculated across all principal components for each time step,

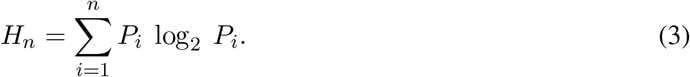

#### Automatic Classification

A linear Support Vector Machine (SVM) method trained was trained and tested using a 5-fold cross-validation which was repeated for one hundred iterations to ensure robustness (toolbox `Classification Learner`, MatLab^™^). The convolved responses of the neurone model output which constituted the input to the algorithm comprised vector data binned into eleven classes representing eleven different characteristic times for stimulation profiles.

### Neurone model

The Izhikevich bifurcating neurone model (74) was used to produce spiking responses. This model includes two state variables, *v*, to represent the membrane potential and, *u*, to represent membrane recovery acting as negative feedback. The dynamics is expressed by a nonlinear system of two coupled ordinary differential equations where *i* stands for the input current,

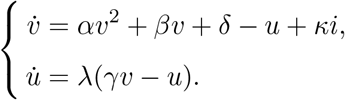

The parameters were selected to approximate biological data (*α* = 0.04, *β* = 5, *δ* = 140, *λ* = 0.02, *γ* = 0.2) (75). The factor κ (10^5^) established the correspondence between mechanoreceptor activation and input current resulting from the opening of ion channels. When the membrane potential, *v*, reached a depolarisation threshold, ξ (+30 mV), a spike was produced accompanied by a reset of the membrane potential at *v_0_* (−65 mV) and an increment of the membrane recovery potential by *u_0_* (+9 mV),

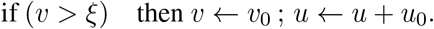

The neurone model is deterministic. To represent the stochasticity of actual neurones (76), Gaussian white noise was added to *v* and *u* in the respective governing equations (function `awgn`, MatLab^™^). To represent the post-synaptic potential, the modelled spike output was convolved with the kernel (77),

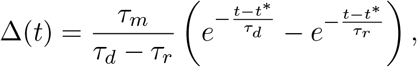

where, *t*\* is the time of spike occurence, *τ_d_* is the decay time (4 ms), *τ_r_* is the rise time (12.5 ms) and *τ_m_* is a normalising time (21.3 ms) to match values from *in vivo* data.

## Data Availability

The full set of mechanical data used to produce all of the results in this paper is available from the corresponding author upon request.

## Acknowledgements

This work was supported by the EU H2020 Grant FETOpen project #829186 ph-coding (Predictive Haptic COding Devices In Next Generation interfaces)

## Contributions

U.B.R., H.J., and V.H. planned and designed the study. U.B.R., H.J., V.H., and A.S wrote the paper; U.B.R. and H.J. performed research; U.B.R., H.J., and V.H. contributed new reagents/analytic tools; U.B.R. and H.J. analysed data.

## Competing Interests

The authors declare that they have no competing interests.

## Correspondence

Correspondence and requests for materials should be addressed to

Udaya Rongala.

## Notes

### Competing Interest Statement

The authors have declared no competing interest.

### Summary of Updates

Abstract, Introduction, Results, Discussion, Methods. New Results were added. The model was adapted.

